# Mathematical modeling reveals a complex network of signaling and apoptosis pathways in the survival of memory plasma cells

**DOI:** 10.1101/2021.08.26.457784

**Authors:** Philipp Burt, Rebecca Cornelis, Gustav Geißler, Stefanie Hahne, Andreas Radbruch, Hyun-Dong Chang, Kevin Thurley

## Abstract

The long-term survival of memory plasma cells is conditional on the signals provided by dedicated survival niches in the bone marrow organized by mesenchymal stromal cells. Recently, we could show that plasma cell survival requires secreted factors such as APRIL and direct contact to stromal cells, which act in concert to activate NF-kB- and PI3K-dependent signaling pathways to prevent cell death. However, the precise dynamics of the underlying regulatory network are confounded by the complexity of potential interaction and cross-regulation pathways. Here, based on flow-cytometric quantification of key signaling proteins in the presence or absence of the required survival signals, we generated a quantitative model of plasma cell survival. Our model emphasizes the non-redundant and essential nature of the two plasma cell survival signals APRIL and stromal cell contact, providing resilience to endoplasmic reticulum stress and mitochondrial stress, respectively. Importantly, the modeling approach allowed us to unify distinct data sets and derive a consistent picture of the intertwined signaling and apoptosis pathways regulating plasma cell survival.

## Introduction

The vertebrate immune system has the unique ability to provide long-term protection against pathogens it has encountered previously. An important cellular correlate of this long-term immunity is the long-lived memory plasma cell which constitutively secretes copious amounts of specific antibodies. Antibodies produced by memory plasma cells serve as a highly efficient barrier against re-infection with pathogens, but have also been associated with a variety of autoimmune diseases (1–3). Memory plasma cell survival is conditional on signals provided to them in dedicated survival niches, organized by mesenchymal stromal cells most prominently in the bone marrow, where memory plasma cells can survive for decades (1,3–6). The survival of memory plasma cells is mediated through the integration of multiple input signals, including integrin-mediated contact to mesenchymal stromal cells and signaling via B cell maturation antigen (BCMA) and Transmembrane activator and CAML interactor (TACI) (7,8). Recently, we demonstrated that provision of A proliferation inducing ligand (APRIL), addressing BMCA, and cell contact to a stromal cell, maintains primary bone marrow memory plasma cells in vitro via NF-kB (nuclear factor ‘kappa-light-chain-enhancer’ of activated B-cells) and PI3K (phosphatidylinositol 3-kinase) signaling pathways, respectively (9). We could show that APRIL and cell contact address different cellular stress pathways, namely the endoplasmic reticulum stress and mitochondrial stress, respectively. However, the architecture of the intracellular signaling network regulating the resilience to these stressors and thereby promoting plasma cell survival remains poorly understood.

A prominent regulatory mechanism restricting the life-span of plasma cells is the widely studied Bcl-2-associated X protein (BAX)-dependent apoptosis pathway (10). Apoptosis is regulated by extrinsic and intrinsic signals and ultimately results in the activation of caspases, a family of cysteine proteases. Mitochondria play a fundamental role in the coordination of the apoptotic pathways. Oligomerization of two proteins, BAX and BAK (BCL2-antagonist/killer 1), that are localized at the mitochondrial outer membrane and cytosol, lead to the formation of the apoptotic pore. Pore formation precedes cytochrome c and apoptosis-inducing factor release into the cytosol and downstream activation of caspases. The BCL-2 family comprises a group of proteins that is critical for the control of apoptosis by regulating the oligomerization of BAX and BAK. Localized mainly at the mitochondria, they can be divided into pro- and anti-apoptotic proteins. The anti-apoptotic proteins (B-cell lymphoma 2, BCL2 and myeloid cell leukemia 1, MCL1) prevent apoptotic pore formation and preserve the integrity of the mitochondrial membrane by binding to BAX and BAK. Pro-apoptotic proteins (Bcl-2-like protein 11, BIM and NOXA), next to BAX and BAK, compete in binding to the anti-apoptotic proteins thereby neutralizing them. Thus, the ratio of pro- to anti-apoptotic proteins plays a decisive role in apoptosis regulation (11). The expression of the anti-apoptotic proteins BCL2 and MCL1 was shown to be upregulated in plasma cells, but while BCL2 seems to be dispensable for the maintenance of memory plasma cells, MCL1 is essential for their survival (12). In multiple myeloma, MCL1 strongly binds to BIM thereby blocking apoptosis (13). In addition to BIM, MCL1 was reported to interact with NOXA (14).

The complexity of the apoptotic network and the design principles leading to the apoptosis decision have been investigated extensively both by experiment and mathematical models. Pioneered by the work of Fussenegger and colleagues, mechanistic differential-equation based models of caspase signaling were introduced to study apoptosis systems (15,16). Subsequent studies focusing on caspase activation in single cells showed that this process is rapid and irreversible (17,18). Many experimental and computational studies have since then successfully demonstrated that the apoptosis decision is well-described by a bi-stable and reversible switch (19–21). Several combined efforts of modeling and experiment have focused on quantitative aspects of the apoptotic network either in large-scale studies or focusing on individual control points of the apoptosis network (22,23). While many of the mechanisms regulating apoptosis have been quantified and modeled, it is less clear how different anti-apoptotic extracellular input signals affect the delicate machinery of cell death, and how well cell-type specific aspects are captured by the available modeling framework.

In this work, we developed a mathematical model of plasma cell survival in the bone marrow, using published (9,24) and yet unpublished data on key components of the plasma cell apoptosis pathway. To this end, we started from a core model based on established principles of the BAX-dependent apoptosis pathway, by adding the extracellular input signals APRIL and ST2 cells, and subsequently, by considering a more detailed interaction network of several caspase proteins downstream of the BAX-module. We found that the survival factors APRIL and ST2 cells have differential rather than additive roles in the regulation of plasma-cell lifespan extension, acting in different ways and primarily on different parts of the network. Further, our analysis underlines the essential aspect of differential caspase regulation for the apoptosis decision and provides insight into the parameters that could be manipulated to alter plasma cell lifespan.

## Methods

### Cell culture and Flow cytometric measurements of apoptosis proteins

Experiments were performed with the same cell culture system as previously reported (9). Intracellular antigens were stained by fixing cells with PFA and permeabilization with methanol. To prevent unspecific binding, cells were pre-incubated with blocking buffer and subsequently stained with primary antibody for 1 hour and, if necessary, with secondary antibody for 30 minutes. Samples were analyzed using a MacsQuant analyzer and FlowJo software. Cytometric procedures followed the recommendations of the “Guidelines for use of flow cytometry and cell sorting in immunological studies” (25). The following antibodies were used in the experiments:

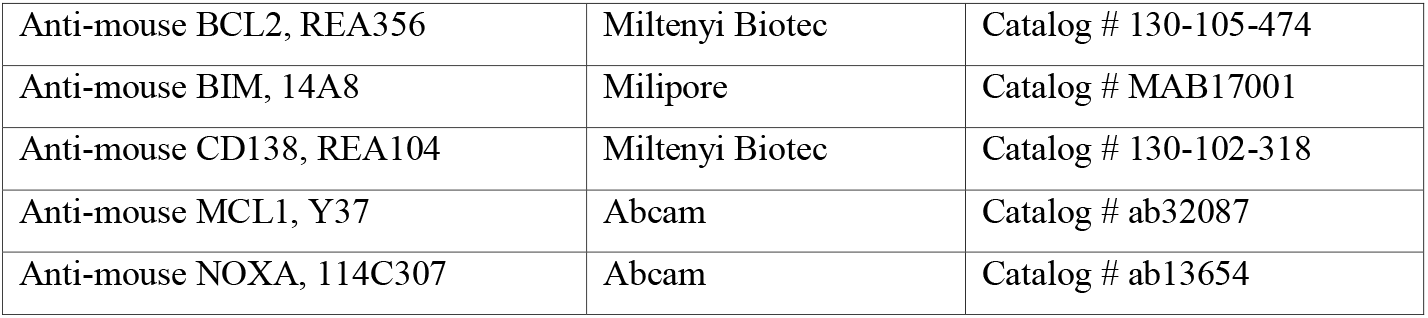

### Data analysis and statistics

To estimate the half-life for plasma cells under different conditions, we fitted an exponential decay function ***f*(*t*) = 100*e*^−*λt*^** to each individual time series. The resulting decay rates were converted to half-lives for each condition according to: ***t*_1/2_ = *log*(2)/*λ***, where ***λ*** represents the average decay rate summarized from the individual fit procedures. To compare average protein concentrations, geometric means for each protein measured under a specific condition were first normalized to the respective medium condition and then compared using an unpaired Student’s t-test. Uncertainties of protein ratios were calculated from the normalized protein concentrations by bootstrapping. To this end, we repeatedly drew from the original samples with replacement to estimate mean and standard deviation of the data.

### Mathematical models and numerical simulations

All model simulations were carried out in Python 3.8. Ordinary differential equations were solved using the scipy.odeint routine. For curve-fitting, least-squares optimization was employed using the Levenberg–Marquardt algorithm implemented in the Python lmfit library. For the perturbation analysis, we defined the effect size as the log2 fold-change of half-life (or BAX*) between a model simulation with best-fit parameter values and the model simulation with perturbed parameters. The parameters were either up- or downregulated by one order of magnitude.

#### a) BAX-dependent apoptosis

We consider (i) production and degradation of MCL-2 family proteins in dependence of the input stimuli APRIL and ST2, (ii) complex formation between pro- and anti-apoptotic proteins (see Figure 2B), and (iii) a dependence of the average half-life of the plasma cell population on the average concentration of activated BAX (BAX*) as follows:

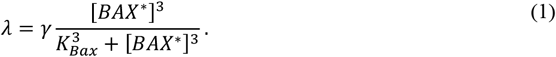

A complete description of the model equations is available as Supplemental Text, parameters are listed in Table 1.

**Table 1:** Parameter values used in the mathematical models.

| Parameter | Value | Unit | Role | Source |
| --- | --- | --- | --- | --- |
| $a_{BCL-2}$ | 0.11 | - | Effect APRIL on $g_{BCL-2}$ | Fit |
| $s_{MCL-1}$ | 0.37 | - | Effect ST2 on $g_{MCL-1}$ | Fit |
| $s_{BCL-2}$ | 0.53 | - | Effect ST2 on $g_{BCL-2}$ | Fit |
| $s_{BIM}$ | 0.49 | - | Effect ST2 on $g_{BIM}$ | Fit |
| $s_{NOXA}$ | 0.40 | - | Effect ST2 on $g_{NOXA}$ | Fit |
| $\gamma$ | 0.43 | $d^{-1}$ | Max. effect size BAX* | Fit |
| $\kappa$ | 1.78 | - | Max. effect Caspase 12 | Fit |
| $\beta$ | 2.65 | - | Inhibition strength ST2 on Caspase 3,7 | Fit |
| $d_{MCL-1}$ | 16.4 | $d^{-1}$ | Decay rate MCL-1 | [30] |
| $d_{BCL-2}$ | 0.86 | $d^{-1}$ | Decay rate BCL-2 | [30] |
| $d_{BIM}$ | 5.94 | $d^{-1}$ | Decay rate BIM | [32] |
| $d_{NOXA}$ | 32.8 | $d^{-1}$ | Decay rate NOXA | [33] |
| $d_{BAX}$ | 1.38 | $d^{-1}$ | Decay rate BAX | [31] |
| $K_{d,3}$ | 2.0 | nM | Dissociation constant | [24] |
| $K_{d,4}$ | 22.0 | nM | Dissociation constant | [24] |
| $K_{d,5}$ | 40.0 | nM | Dissociation constant | [24] |
| $K_{d,6}$ | 2.50 | nM | Dissociation constant | [24] |
| $K_{d,7}$ | 68.0 | nM | Dissociation constant | [24] |
| $k^+$ | 0.17 | $\mu M d^{-1}$ | Complex association rate | - |
| $k_1$ | 43.2 | $\mu M d^{-1}$ | BAX activation rate | - |
| $k_2$ | 8.64 | $\mu M d^{-1}$ | BAX* deactivation rate | - |
| $g_{p,0}$ | 0.86 | $\mu M d^{-1}$ | Basal protein growth | - |
| $\alpha$ | 10.0 | - | Inhibition strength APRIL on Caspase 12 | - |
| $K_{BAX}$ | 200 | nM | Half-saturation constant BAX-halflife | - |

#### b) BAX-independent regulation of caspases

To consider the effect of direct caspase regulation by APRIL and ST2 on the apoptosis decision, we extended the BAX-Apoptosis model and assumed additional regulation of the cell-death rate λ as follows (see Figure 3A and text):

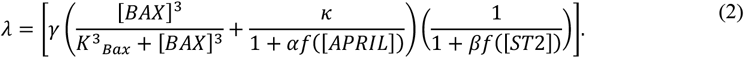

That means, the cell-death rate *γ* is increased due to the activity of caspase 3/7, which is induced through a combination of BAX-dependent and independent effects. Those BAX-independent effects stem from activity of caspase 12, which can be inhibited by APRIL. Finally, the activity of caspase 3/7 can be inhibited by ST2. Here, the fitting parameter *κ* is the relative effect of caspase 12 activity, the fitting parameters β denotes the maximal inhibitory effect ST2 on caspase 3/7, and we set *α* = 10 for the inhibitory effect size of APRIL. For model fitting to our data in absence and presence of APRIL and ST2, we adopt a Boolean formulation for the regulatory function, 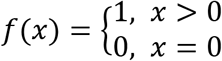. Modification of this function into a more specific Hill-type form such as *f*(*x*) = *x^n^*/(*x^n^* + *K*) is straight-forward.

## Results

### A quantitative model of BAX-dependent apoptosis in plasma cells

Previous experiments have shown that plasma cells are not intrinsically long-lived but that provision of the cytokine APRIL combined with cell-cell contact, conferred by the stromal cell line ST2, keeps them alive in vitro, while each factor alone proved to be insufficient (9)(Figure 1A). Indeed, re-plotting the data of (8) separately for individual experiments revealed an exponential survival curve for cultured plasma cells, where the half-life is significantly increased from less than 1 day to more than 6 days after addition of APRIL and ST2 (Figure 1B-C and Figure S1). How these signals integrate and affect the underlying complex regulatory network facilitating apoptosis decisions remains unclear (Figure 1A). In particular, we wondered whether the experimental observations could be reconciled with an additive model, where APRIL and ST2 synergistically act to inhibit BAX-dependent apoptosis, or alternatively, whether APRIL and ST2 address fundamentally distinct parts of the apoptosis network.

**Figure 1:**
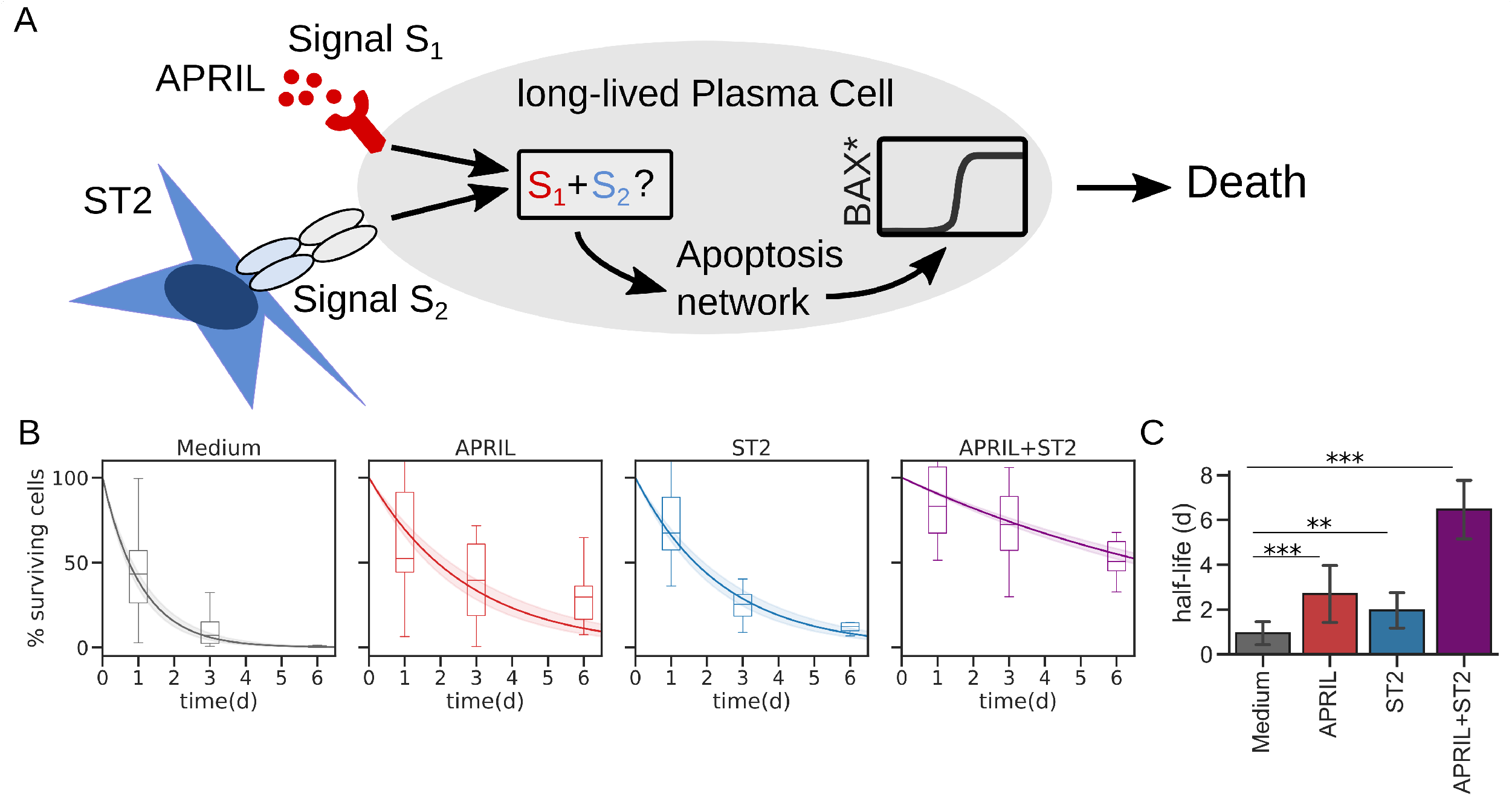
Regulation of plasma cell survival via APRIL and ST2. (A) Schematic signal integration of ST2 and APRIL. How the signals are integrated into the Bax-dependent apoptosis network is yet unclear. Time course were data taken from Ref. (9) and are here represented as Box plots. Data were fitted to exponential curves, each experiment at a time (Figure S1). Curves shown here represent mean+s.e.m. of those fit results. C) Average half-lives of each condition based on the fitting procedure from (B). **P<0.01, ***P<0.001, n>12 fits per condition. Error bars represent standard deviation.

Since in plasma cells, APRIL stimulates the NF-kB signaling pathway (26–28)(8) and ST2 acts via the PI3K pathway to stimulate the transcription factor FoxO (9,29), we started our analysis based on the working-hypothesis that ST2 and APRIL act by targeting components of the BAX-dependent apoptosis pathway. To derive a specific mathematical model, we measured the abundance of several key components of the BCL-2 family, namely BIM, BCL-2, NOXA and MCL-1, after 3 days in our established in vitro culture system (9) with and without stimulation by APRIL and ST2 by immunofluorescent single-cell staining (Figure 2A). We found that the presence of APRIL alone had a tendency to affect BCL-2 (p=0.123), whereas ST2 had a significant negative effect on BIM, NOXA and MCL-1 concentrations (Figure 2A). The divergent effects of APRIL and ST-2 on different members of the BCL-2 family were even more pronounced when considering the ratios within pairs of pro- and anti-apoptotic proteins (Figure 2B). We used that information to establish a refined regulatory network of BAX-dependent plasma-cell apoptosis (Figure 2C). Combined with a recently published compilation of quantitative data for the BCL-2 interactome (Table 1)(24,30–33), we were now in a position to formulate and annotate a specific mathematical model of BAX-dependent apoptosis in plasma cells (Methods and Supplementary Text). The only parameters lacking good estimates from the literature are the values describing protein production rates, which we therefore used as fitting parameters. Indeed, the resulting model was able to describe the new data set of BCL-2 family member protein abundance (Figure 2D).

**Figure 2:**
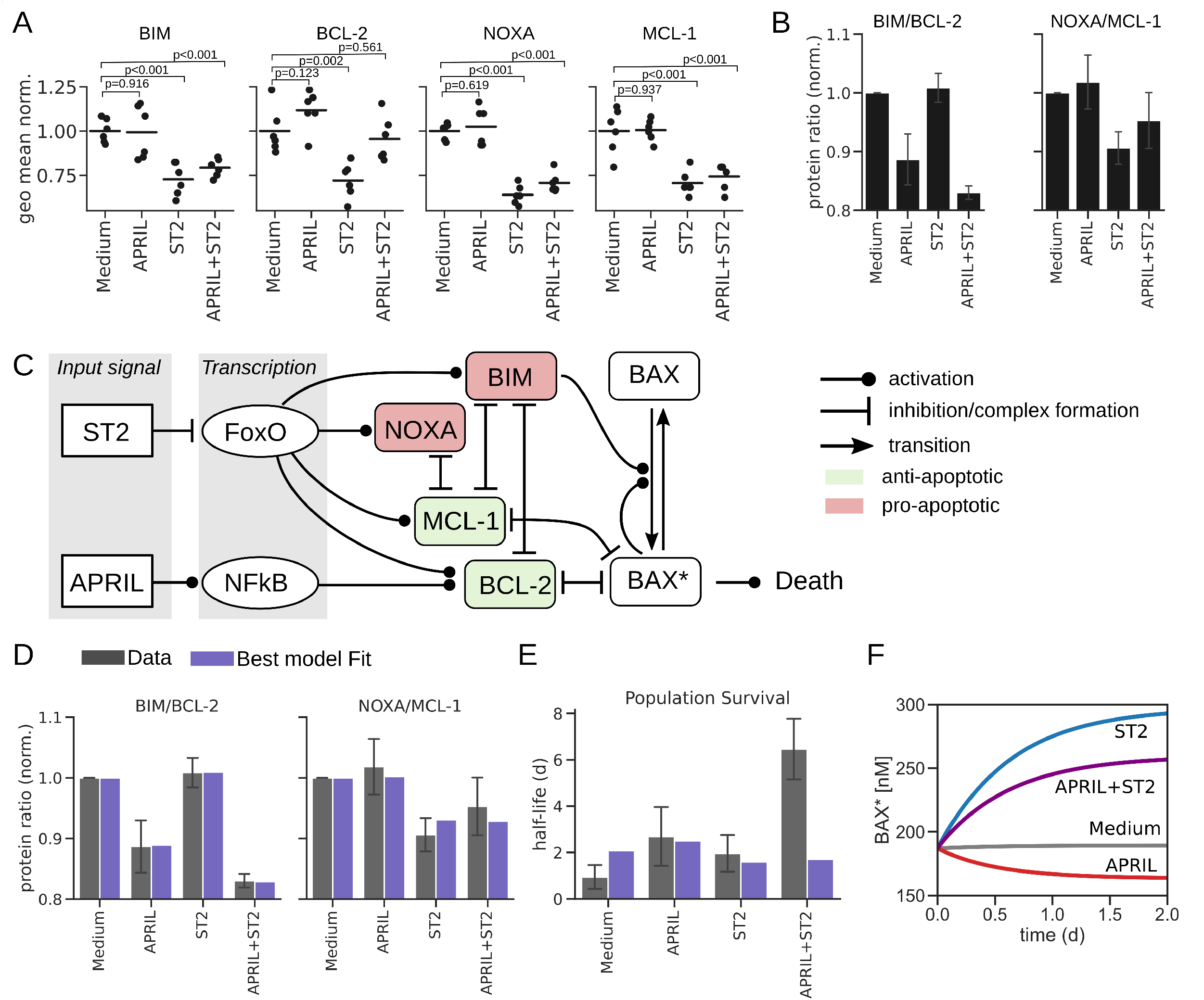
A mechanistic model quantifies the BAX-dependent apoptosis network in plasma cells. (A) Geometric mean of measured core proteins in APRIL/ST2/APRIL+ST2 environments normalized to the respective concentration without stimulus (Medium). (B) Ratios of indicated proteins taken from (A) after normalization to Medium condition. (C) Model scheme based on the effects of APRIL and ST2 on MCL-2 family protein abundance data shown in A. The model includes pro-apoptotic proteins BIM and NOXA (red) and anti-apoptotic proteins BCL-2 and MCL-1 (green). (D) Protein ratios (see A) fitted to the model shown in panel B. Error bars represent standard deviation. (E) Model fit to survival kinetics (Figure 1B) after fitting the protein ratios. (F) Time-course simulation of the mechanistic apoptosis model for different inputs as indicated.

Having established a mathematical formulation of the BAX-dependent apoptosis pathway in the context of plasma cell survival, we proceeded to the original question of whether APRIL and ST2-dependent regulation of that pathway are sufficient to explain the observed survival kinetics shown in Figure 1B. Interestingly, we found that our model was well able to fit the protein and survival data individually (Figure S2), but it was not able to capture both protein and survival data at the same time within the expected range of the experimental data (Figure 2E). In fact, the annotated model predicted a positive regulatory effect of ST2 on the average level of activated BAX (BAX*) in the plasma cell population (Figure 2F). That ST2-dependent elevation of BAX* cannot be fully compensated for by APRIL, which explains the disconnect of that data-derived model parameterization with our cell-survival data (Figure 2E). Hence, we concluded that APRIL and ST2-driven regulation of BAX-dependent apoptosis alone is insufficient to explain the observed survival curves, and rather, additional regulatory mechanisms must be considered.

### Direct regulation of Caspase proteins is required for effective control

Caspases are not only mediators, but also critical regulators of apoptosis (34,35), and our recent data suggest that APRIL and ST2 have different roles in their regulation (9). Specifically, we found that ST2 inhibits caspases 3 and 7, whereas APRIL inhibits the ER-stress-induced apoptosis mediated via caspase 12. Therefore, we supplemented our core model of BAX-dependent apoptosis by considering a network of caspase regulation (Figure 3A)(Methods). To specify the model, we added the 2 new fitting parameters describing the extended caspase network, and we found that this extended model indeed captures the individual and combined effects of APRIL and ST2 (Figure 3B-C). In particular, in contrast to the BAX core model, we obtained good results fitting the new caspase-related parameters to the plasma cell survival data (Figure 3C).

**Figure 3:**
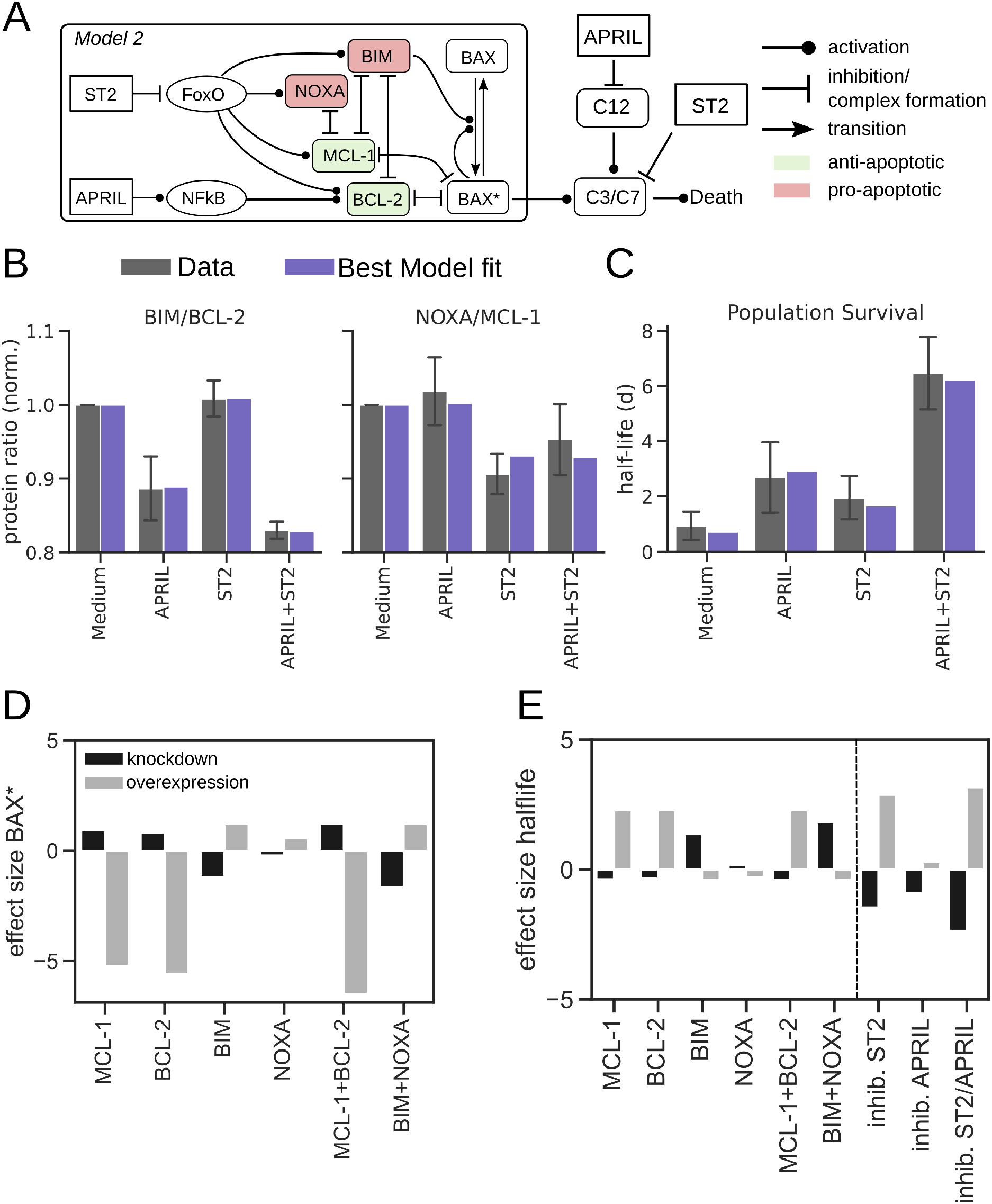
Unifying the data sets on MCL-2 family members and plasma cell survival data requires direct regulation of caspases. (A) Model scheme, combination of the mechanistic mitochondrial apoptosis model (see Figure 2B) with ER-stress-induced caspase activation. (B-C) Combined model fit to protein data and survival kinetics. Error bars represent standard deviation. (D-E) Effect on half-life and BAX activation for simulated knockdown or overexpression of proteins in the apoptosis network. Growth rates for protein species were varied by one order of magnitude in both directions.

The successful data annotation gave us the opportunity to analyze the dependency on individual stimuli and on different parts of the pathway in more detail. Interestingly, we found that in the model, overexpression of anti-apoptotic proteins is far more effective than knock-down of pro-apoptotic proteins in the regulation of activated BAX (Figure 3D). Further, regulation of caspase 3/7 turned out to have the largest effect on plasma cell life-span (Figure 3E). That is in good agreement with the intuition gained from Figure 2E, that direct regulation of caspase 3/7 through ST2 is required to counteract the pro-apoptotic role of ST2 within the BAX-dependent apoptosis pathway. Taken together, our analysis suggests that a unified description of the two considered data sets requires consideration of both BAX-dependent and BAX-independent regulation of caspases via both APRIL and ST2.

### Full model topology is essential to describe plasma cell survival

To further test whether our proposed model topology was necessary to describe all available data, we derived a set of sub-models lacking one or more components of the full model (Figure 4A). For each possible combination, we fitted the model to the available data and compared the fitting result to the original model fit (Figure 4B and Figure S3). The full model had clearly the smallest fitting error (*χ*^2^), and models lacking regulatory effects of either APRIL, ST2 or both on caspase activity all provided similar fitting quality (Figure 4B). For a refined model comparison, we employed Akaike’s information criterion (AIC) (Figure 4C), a well-established metric for comparing models with differing numbers of fitting parameters (36). AIC values lack a direct interpretation, and thus we only consider differences between models (-ΔAIC), here we show -ΔAIC in relation to the model with poorest fit quality (Figure 4C). A difference of ΔAIC>2 is usually regarded as significant, and therefore, our analysis clearly rules out all sub-models (Figure 4C). Nevertheless, the ΔAIC representation can also be regarded as a ranking of the most critical model components for explanation of the available data. As such, it is intriguing that in our model, ST2-driven regulation of caspase 3/7 is the most critical individual factor (panel (iv) in Figure 4A and C), even exceeding the effect of by-passing caspase regulation completely (panel (vi)). Hence, our model analysis supports both a strong role for BAX-independent caspase regulation and the critical need of ST2-derived signals for effective regulation of plasma cell survival.

**Figure 4:**
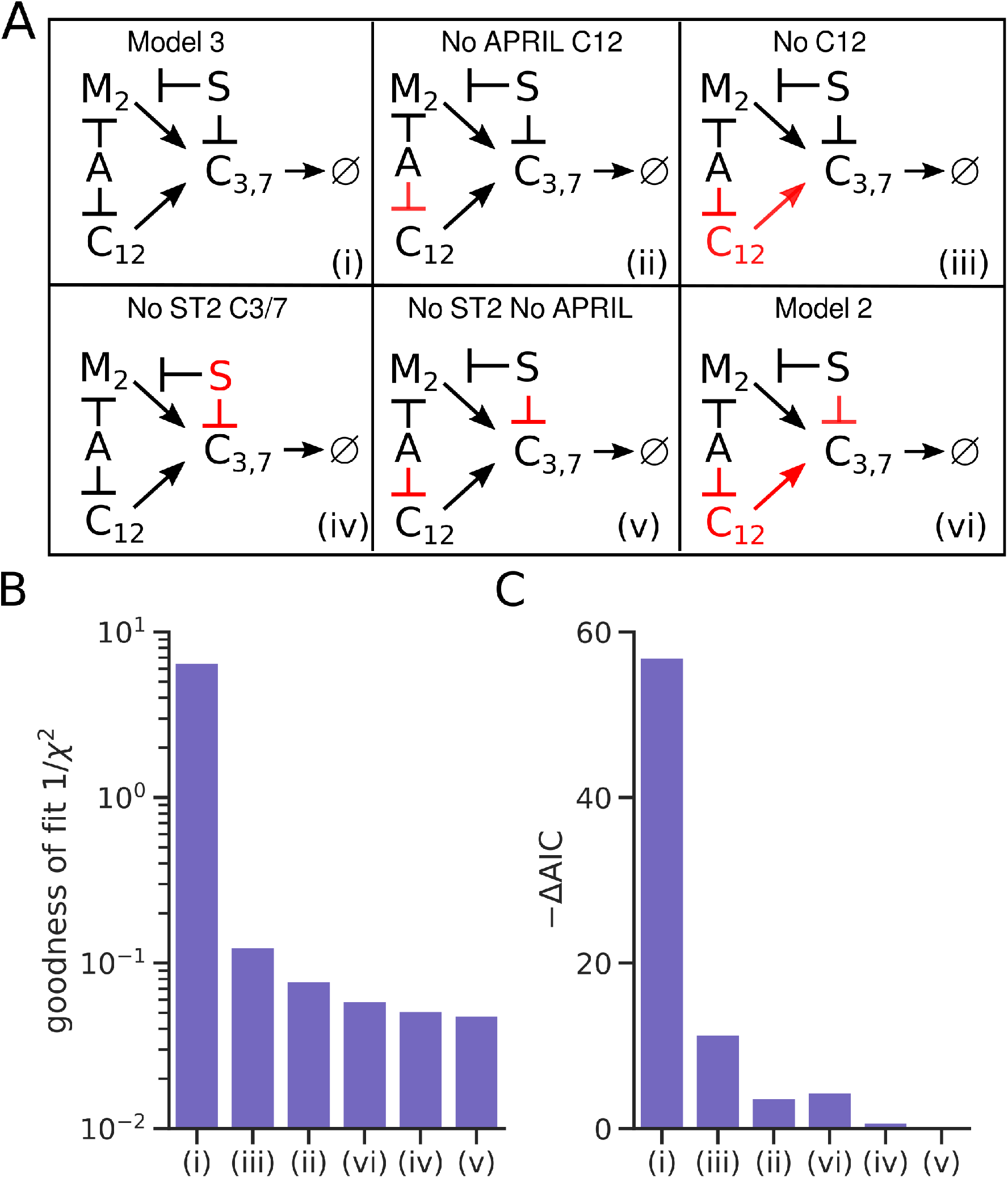
All considered regulatory processes are required to explain the data. (A) Different hypothetic network topologies (ii-vi) as submodules of the model shown in (3A), here presented as a condensed version (i). Abbreviations: M2: Model 2; A: APRIL; S:ST2; C12: Caspase 12; C3/7: Caspase 3, Caspase 7. Model components removed from the full model are shown in red. (B) goodness-of-fit (1/root-mean-squared error)) for each submodel (ii-vi). (C) Akaike information criterion (-ΔAIC) for each sub-model compared to the full model (i), where the model with smallest -ΔAIC is used as a reference and set to 0.

## Discussion

In this work, we developed a mathematical model describing the survival of plasma cells in the bone marrow. We used the model to unify two acquired data sets, namely abundance of pro- and anti-apoptotic proteins and plasma-cell survival kinetics in the presence of the soluble factor APRIL and/or stromal ST2 cells as cell-contact dependent survival signal. In the annotated model, the balance of survival proteins is a critical factor for the lifespan of plasma cells. Further, we found that a combination of direct caspase regulation together with regulation of BAX-dependent pathways is essential to explain all available data.

A special feature in plasma cell biology is the contribution of the endoplasmic reticulum (ER) to apoptosis regulation (10,37). Since plasma cells as the primary antibody-secreting cell type produce vast amounts of proteins, they experience high ER-stress due to protein misfolding. In plasma cells, BAX was not only found in the mitochondrial membrane but also localized at the ER (10). We could recently show that the plasma cell microenvironment, composed of stromal cell contact and the cytokine APRIL, counteracts the activation of caspases 3 and 7 and caspase 12, respectively. Caspases 3 and 7, that are activated upon mitochondrial stress (34), are regulated by PI3K signaling that is activated by cell contact to stromal cells. Activation of the ER-associated caspase 12 is inhibited by APRIL signaling via the NF-kB pathway (9). Here, our model simulations supported the view that differential regulation of caspases by APRIL and ST2 is a key process in the regulation of plasma cell apoptosis.

In vivo, memory plasma cells persist for a lifetime, i.e. they do not have a dedicated half-life: data indicate that plasma cells survive as long as they are provided with their niche (38–40). Our in vitro culturing system cannot yet reflect such very long life-times, which we think is mainly due to technical limitations—the co-cultured ST2 cells start overgrowing the plasma cells after ~5 days in culture. Also, we cannot exclude additional regulatory factors accounting for the very long life-times reported in vivo. However, the observed time-scale separation between a half-life of ~1 day for medium condition and of 6 days in presence of APRIL and ST2 rather argues against the same apoptosis mechanism driving the fast and the slow plasma cell decline. Therefore, in model development, we considered the APRIL+ST2 condition as a base-line reflecting properties of the cell culturing system distinct from apoptosis regulation in the context of long-lived plasma cells.

The life-span of immune cell populations is tightly controlled and is a critical property in a range of inflammatory processes, as shown previously in the context of selective expansion of lymphocyte subtypes (41,42) and NK cell subpopulations (43), amongst others. For therapy, targeted depletion of memory plasma cells has been proposed as a promising strategy in autoimmune diseases such as lupus erythematosus (2). On the other hand, memory plasma cells can provide long-lasting protection against infections, as recently demonstrated for mild SARS-Cov2 infections in humans (44). In plasma cell biology, quantitative studies have previously focused on the generation and composition of the survival niche (45) and on the long-term development of the pool of available memory plasma cells (46), but our quantitative understanding of apoptosis signaling pathways in plasma cells is still quite limited. Here, using cultured plasma cells as a model system, we developed a mathematical modeling framework to study apoptosis regulation in specific immune cell populations in the context of established signaling pathways.

## Funding

This work was supported by the Leibniz association (Junior Research Group program, to K.T.), the Leibniz ScienceCampus Chronic inflammation, the Deutsche Forschungsgemeinschaft (TH 1861/4-1, to K.T. and SFB TRR130 P16 to H.D.C. and A.R.) and the Innovative Medicines Initiative 2 Joint Undertaking under grant agreement no. 777357 to H.D.C. and A.R. H.D.C. is supported by the Dr. Rolf M. Schwiete Foundation.

## Supplementary Material

The supplementary material contains 3 Figures and the detailed description and equations of the BAX-dependent apoptosis model.

## Notes

### Competing Interest Statement

The authors have declared no competing interest.

