## Supplementary Information for "Mathematical modeling reveals a complex network of signaling and apoptosis pathways in the survival of memory plasma cells"

### Supplementary Information - Data-driven modeling reveals a complex network of signaling and apoptosis pathways in the survival of memory plasma cells

#### Authors

Philipp Burt, Rebecca Cornelis, Gustav Geissler, Andreas Radbruch, Hyun-Dong Chang, Kevin Thurley

#### Supplementary Figures

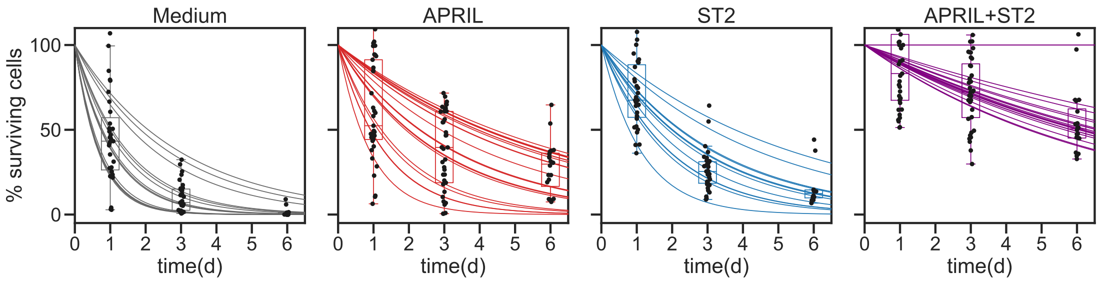


Figure S1: Plasma cell survival kinetics for individual experiments. Each dot represents one measurement for the number of surviving cells for a given time and treatment. Each line (Grey: Medium; Red: APRIL, Blue: ST2, Purple:APRIL+ST2) represents an exponential fit for one experimental series consisting of measurements at three consecutive timepoints (d1, d3, d6).

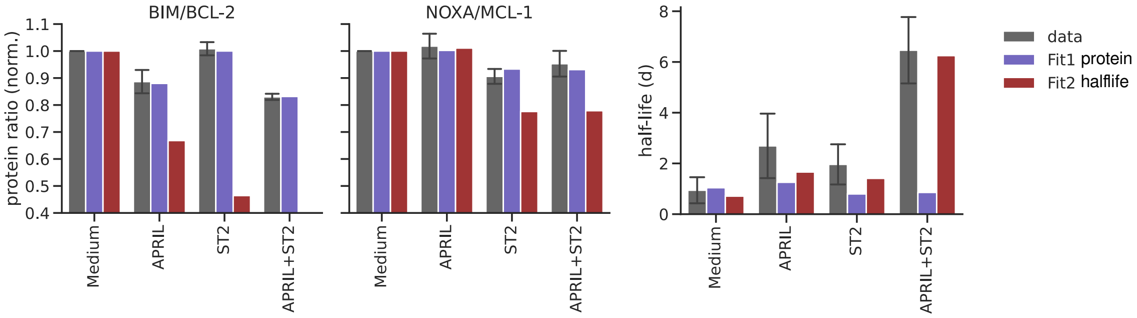


Figure S2: Protein ratio and halflife prediction give good fit results for individual fits but not for both fits combined. Blue and red bars represent model fits with initial conditions that lead to good fit results for either protein ratios (blue bars) or halflives (red bars). Available data for protein ratios and estimated halflives shown in grey. Error bars represent standard deviation.

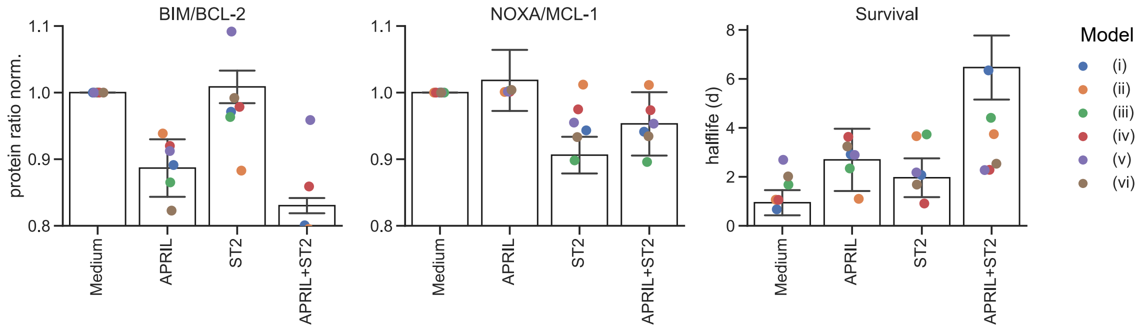


Figure S3: Model fits for protein ratio and halflive based on for different model versions as discussed in Fig 4, main text.

#### Supplementary Text

##### Modeling BAX activation

Focusing on the available data of apoptotic regulators in plasma cells, we developed an ordinary differential equations model based on mass-action-kinetics. The crucial player in our model is the pore-forming protein BAX, which upon activation facilitates the formation of the apoptotic pore. This membrane permeabilization in turn leads to the release of Cytochrome C and subsequently to the activation of downstream caspases and cell death [1, 2, 3]. For BAX activation we consider the following reactions:

$$\text{BIM} + \text{BAX} \xrightarrow{k_1} \text{BIM} + \text{BAX*} \tag{1}$$

$$\text{BAX*} + \text{BAX} \xrightarrow{k_1} 2\text{BAX*} \tag{2}$$

$$\text{BAX*} \xrightarrow{k_2} \text{BAX} \tag{3}$$

With $k_1$ and $k_2$ denoting BAX activation and deactivation respectively. Additionally, we considered here direct activation of BAX via the proapoptotic protein BIM [4]. For further regulation of BAX activation, we considered antiapoptotic proteins BCL-2 and MCL-1 that have been shown to directly bind and form complexes with BAX, thereby inhibiting apoptosis [5]. Moreover, we considered pro-apoptotic proteins NOXA and BIM that have been shown to form heterodimers with anti-apoptotic proteins. Hence, the proapoptotic proteins prevent BAX deactivation by inhibiting the antipoptotic proteins [5]. For complex formation, we therefore consider:

$$\text{MCL-1} + \text{BIM} \underset{k_3^-}{\overset{k^+}{\rightleftharpoons}} \text{MCL-1:BIM} \tag{4}$$

$$\text{MCL-1} + \text{NOXA} \underset{k_4^-}{\overset{k^+}{\rightleftharpoons}} \text{MCL-1:NOXA} \tag{5}$$

$$\text{MCL-1} + \text{BAX*} \underset{k_5^-}{\overset{k^+}{\rightleftharpoons}} \text{MCL-1:BAX*} \tag{6}$$

$$\text{BCL-2} + \text{BIM} \underset{k_6^-}{\overset{k^+}{\rightleftharpoons}} \text{BCL-2:BIM} \tag{7}$$

$$\text{BCL-2} + \text{BAX*} \underset{k_7^-}{\overset{k^+}{\rightleftharpoons}} \text{BCL-2:BAX*} \tag{8}$$

Here $k^+$ and $k_i^-$ denote the complex association and dissociation rate respectively.

**Modeling input of APRIL and ST2 on BAX activation**

We assumed that input signals APRIL and ST2 affect the steady-state concentration of pro- and anti-apoptotic proteins which in turn modulate the availability of activated BAX. Specifically, we assumed that each protein in the network is subject to growth and decay in the simple form

$$\frac{dx_P}{dt} = g_P(\text{APRIL,ST2}) - d_P x_P \tag{9}$$

where APRIL and ST2 affect the growth rate $g_P$. We modeled the growth rate dependency of the inputs as:

$$g_{\text{P}} = g_{\text{P},0}\Big(1 + a_P \frac{[\text{APRIL}]^3}{[\text{APRIL}]^3 + 0.1^3} - s_P \frac{[\text{APRIL}]^3}{[\text{APRIL}]^3 + 0.1^3}\Big) \tag{10}$$

Here, $a_P$ and $s_P$ are the maximal effects of APRIL and ST2 on the growth rate. Since it has been previously shown that ST2 inhibits FoxO, we assume an inhibitory effect for ST2. As described in the main text, we designed the network based on our observations that APRIL only affected BCL-2 whereas ST2 affected BIM, NOXA and MCL-1. Further, we modeled reversible formation of heterodimeric complexes according to:

$$\frac{d[XY]}{dt} = k^+[X][Y] - k^-[XY] - (d_X + d_Y)[XY] \tag{11}$$

Here, $k^+$ is the reaction rate with which the complex is formed and $k^-$ is the reaction rate with which the complex dissociates. For the protein species in our model, quantitative estimates can be mainly found for the dissociation constant $K_D$. Therefore, we chose to set the complex formation rate $k^+$ to a fixed value (see table 1, main text) and calculated the complex dissocation rate as

$$k_i^- = K_{D,i} \cdot k^+ \tag{12}$$

Taken together, this results in the following set of ordinary differential equations (Abbreviations: A: MCL-1; B: BCL-2; C: BIM; D: NOXA; X: BAX; Z: BAX*):

$$\frac{dA}{dt} = g_A - d_A[A] - k^+[A][C] - k^+[A][Z] - k^+[A][D] + k_3^-[AC] + k_5^-[AZ] + k_4^-[AD] \tag{13}$$

$$\frac{dB}{dt} = g_B - d_B[B] - k^+[B][C] - k^+[B][Z] + k_6^-[BC] + k_7^-[BZ] \tag{14}$$

$$\frac{dC}{dt} = g_C - d_C[C] - k^+[A][C] - k^+[B][C] + k_3^-[AC] + k_6^-[BC] \tag{15}$$

$$\frac{dD}{dt} = g_D - d_D[D] - k^+[A][D] + k_4^-[AD] \tag{16}$$

$$\frac{dX}{dt} = g_X - d_X[X] - k_1([C] + [Z])[X] + k_2[Z] \tag{17}$$

$$\frac{dZ}{dt} = -d_X[Z] + k_1([C] + [Z])[X] - k_2[Z] - k^+[A][Z] - k^+[B][Z] + k_5^-[AZ] + k_7^-[BZ] \tag{18}$$

$$\frac{d[AC]}{dt} = -(d_A + d_C)[AC] + k^+[A][C] - k_3^-[AC] \tag{19}$$

$$\frac{d[AD]}{dt} = -(d_A + d_D)[AD] + k^+[A][D] - k_4^-[AD] \tag{20}$$

$$\frac{d[AZ]}{dt} = -(d_A + d_Z)[AZ] + k^+[A][Z] - k_5^-[AZ] \tag{21}$$

$$\frac{d[BC]}{dt} = -(d_B + d_C)[BC] + k^+[B][C] - k_6^-[BC] \tag{22}$$

$$\frac{d[BZ]}{dt} = -(d_B + d_Z)[BZ] + k^+[B][Z] - k_7^-[BZ] \tag{23}$$
